# The formin Fmn2 is required for the development of an excitatory interneuron module in the zebrafish acoustic startle circuit

**DOI:** 10.1101/2020.07.23.218032

**Authors:** Dhriti Nagar, Tomin K James, Ratnakar Mishra, Shrobona Guha, Aurnab Ghose

## Abstract

The formin family member, Fmn2, is a neuronally enriched cytoskeletal remodelling protein conserved across vertebrates. Recent studies have implicated Fmn2 in neurodevelopmental disorders, including sensory processing dysfunction and intellectual disability in humans. Cellular characterization of Fmn2 in primary neuronal cultures has identified its function in the regulation of cell-substrate adhesion and consequently growth cone translocation. However, the role of Fmn2 in the development of neural circuits *in vivo*, and its impact on associated behaviours have not been tested.

Using automated analysis of behaviour and systematic investigation of the associated circuitry, we uncover the role of Fmn2 in zebrafish neural circuit development. As reported in other vertebrates, the zebrafish ortholog of Fmn2 is also enriched in the developing zebrafish nervous system. We find that Fmn2 is required for the development of an excitatory interneuron pathway, the spiral fiber neuron, which is an essential circuit component in the regulation of the Mauthner cell-mediated acoustic startle response. Consistent with the loss of the spiral fiber neurons tracts, high-speed video recording revealed a reduction in the short latency escape events while responsiveness to the stimuli was unaffected.

Taken together, this study provides evidence for a circuit-specific requirement of Fmn2 in eliciting an essential behaviour in zebrafish. Our findings underscore the importance of Fmn2 in neural development across vertebrate lineages and highlight zebrafish models in understanding neurodevelopmental disorders.

**SIGNIFICANCE STATEMENT:** Fmn2 is a neuronally enriched cytoskeletal remodelling protein linked to neurodevelopment and cognitive disorders in humans. Recent reports have characterized its function in growth cone motility and chemotaxis in cultured primary neurons. However, the role of Fmn2 in the development of neural circuits *in vivo* and its implications in associated behaviours remain unexplored. This study shows that Fmn2 is required for the development of neuronal processes in the acoustic startle circuit to ensure robust escape responses to aversive stimuli in zebrafish. Our study underscores the crucial role of the non-diaphanous formin, Fmn2, in establishing neuronal connectivity and related behaviour in zebrafish.

## INTRODUCTION

The behavioural repertoire of an organism is contingent on accurate neural connectivity of the nervous system. During the development, individual neurites are guided to reach their predetermined targets by environmental cues. Specialized tips of growing neurites, the growth cones, employ a plethora of cytoskeleton regulatory proteins to locally remodel the cytoskeleton and execute directional translocation in response to guidance cues (Lowery and Van Vactor, 2009). Precisely regulated, dynamic remodelling of the actin and microtubule cytoskeletons enables the growth cone to navigate towards its synaptic target accurately. Actin rich growth cone filopodia express guidance receptors and function as chemosensory antennae. Differential stabilization of filopodia, via microtubule capture, allows the microtubule network to advance in a spatially biased manner to bring about directional forward displacement (Dent et al., 2011). Further, the cytoskeleton modulates the adhesion of the growth cone to the surrounding matrix and the ability to generate traction forces (Kerstein et al., 2015). There are several classes of proteins involved in active cytoskeleton remodelling. One such family of proteins is the formin family of proteins. Formins are a conserved family of actin nucleators and processive elongators of non-branched actin filaments characterized by the FH1 and FH2 domains (Higgs, 2005; Higgs and Peterson, 2005; Breitsprecher and Goode, 2013). Members of the formin family are known to regulate other aspects of cytoskeleton dynamics, like F-actin bundling, actin-microtubule crosstalk and the adhesion of the growth cone to the extracellular matrix (Kawabata Galbraith and Kengaku, 2019).

Most formins are broadly expressed in multiple tissues types (Schönichen and Geyer, 2010; Breitsprecher and Goode, 2013; Krainer et al., 2013; Dutta and Maiti, 2015; Kawabata Galbraith and Kengaku, 2019) but the non-diaphanous related formin, Formin-2 (Fmn2), is found to be enriched in the developing and the mature nervous systems of mice, humans and chicken (Leader and Leder, 2000; Katoh and Katoh, 2004; Dutta and Maiti, 2015; Sahasrabudhe et al., 2015). Fmn2 has been implicated in neurodevelopmental disorders, intellectual disability, age-related dementia, microcephaly and sensory processing dysfunction in humans (Perrone et al., 2012; Almuqbil et al., 2013; Law et al., 2014; Agís-Balboa et al., 2017; Anazi et al., 2017; Marco et al., 2018). Recent studies identify the involvement of Fmn2 in regulating of spine density in hippocampal neurons (Law et al., 2014) and in the development of the corpus callosum in humans (Gorukmez et al., 2020).

So far, there is little known about the mechanisms by which Fmn2 causes neurodevelopmental defects. Fmn2 is required for growth cone motility and axonal pathfinding of spinal neurons. Filopodial stability and the dynamics of the adhesion complexes between the growth cone and the substrate are regulated by Fmn2 (Sahasrabudhe et al., 2016; Ghate et al., 2020). Another study has identified actin-microtubule crosstalk mediated by Fmn2 as a mechanism underlying growth cone turning (Kundu et al., 2020). These reports primarily use cultured primary neurons which precludes investigating gene function in assembling neural circuits giving rise to appropriate behaviours.

Zebrafish is an excellent vertebrate model to interrogate gene function in the establishment of neural circuits *in vivo* and associated behaviours (Kalueff et al., 2013; Mcarthur et al., 2020). One of the most studied neural circuits is the Mauthner cell mediated acoustic startle circuit (Swain et al., 1993; Jontes et al., 2000; Korn and Faber, 2005; Burgess et al., 2009; Sillar, 2009; Issa et al., 2011; Kinkhabwala et al., 2011; Hale et al., 2016; Liu and Hale, 2017; Hecker et al., 2020). Acoustic and tactile stimuli from the environment are processed by the M-cells aided by regulatory excitatory and inhibitory interneuron module and relayed to the motor neurons downstream (Korn and Faber, 2005). Mauthner mediated escapes are essential to the survival of the zebrafish in the wild environment. The acoustic startle circuit comprising of sensory inputs to the M-cells and regulatory interneurons imparts the ability to respond in time to evade a predator.

In this study, we show that the Fmn2 expression is enriched in the zebrafish nervous system and is necessary for the development of an excitatory interneuron pathway indispensable for Mauthner cell-mediated fast escape responses to aversive stimuli.

## MATERIALS AND METHODS

### Zebrafish maintenance

Locally sourced wildtype strain of zebrafish, bred in the lab for three generations were used in all the experiments. Adult zebrafish were raised in a recirculating aquarium system (Techniplast) at 28.5°C under a 14-hour light and 10-hour dark cycle. Embryos were collected and raised in E3 buffer (5 mM NaCl, 0.33 mM MgSO_4_, 0.17 mM KCl, 0.33 mM CaCl_2_, 5% Methylene Blue) at 28.5°C and used at different stages as previously described (Kimmel et al., 1995). The following transgenic zebrafish lines were used.

Tg(Cldn-b:Lyn-GFP) (Haas and Gilmour, 2006)

Tg(Brn3c:GAP43-GFP) (Xiao et al., 2005)

### Whole mount *in situ* hybridization

Total RNA was isolated from 48 hours post-fertilization (hpf) wildtype embryos using the RNeasy mini kit (Qiagen) and reverse transcribed using the SuperScript IV RT Kit (ThermoFisher) to obtain cDNA. The cDNA was amplified for a 366 bp long gene specific region of Fmn2 corresponding the 5’ UTR and exon 1 flanked with T7 and T3 promoter sequences in the antisense and sense direction respectively.

Primer sequences:

Fmn2_ISH_5UTR_F_T3:

GCAATTAACCCTCACTAAAGGGATGCGTTGTTGTGTTTGTG

Fmn2_ISH_5UTR_R_T7:

TAATACGACTCACTATAGGGGCTCTCGCTGTCTGATGAAG

The amplified product was purified and used as the template for in vitro transcription of antisense and sense probes against Fmn2 mRNA. Zebrafish embryos ranging from 1 cell stage to 96 hpf were used for whole-mount in situ hybridization experiments as described previously (Thisse and Thisse, 2008). BM Purple was used as a chromogenic substrate for detection.

### Morpholino and RNA injections

Two morpholinos targeting Fmn2 were obtained from Gene Tools. The splice blocking morpholino targets the intron between exons 5 and 6 and ensures the inclusion of a stop codon in the translation frame. The translation blocking morpholino binds early on in the first exon. The sequences of the morpholinos used are given below.

MO Control: 5’-CCTCTTACCTCAGTTACAATTTATA-3’

MO SB Fmn2: 5’-ACAGAAGCGGTCATTACTTTTTGGT-3’

MO TB Fmn2: 5’-ATGAGCGGCGGCGGTTTCAAGCCAT-3’

All experiments were done by injecting 2 nl volume of the MO Control and MO SB Fmn2 (1 ng/nl; 2 ng per embryo) in the cytoplasm of the 1-cell stage embryos. For MO TB Fmn2 dose dependence, 4 ng/embryo and 8 ng/embryo doses of the morpholino were injected in the yolk of embryos in addition to the cytoplasmic injection of 2 ng/embryo. The injection volume was calibrated at 2 nl per embryo for each needle before injection. Post injections, embryos were washed and raised in E3 buffer (supplemented with methylene blue) at 28.5 °C till the desired developmental stage with regular cleaning. For immunostaining and live imaging experiments, the buffer was supplemented with 0.003% Phenylthiourea (PTU; Sigma) to remove pigmentation from the skin. For rescue experiments, capped mRNA was synthesized using the SP6 mMessage mMachine RNA synthesis kit (Ambion) from pCS2-mFmn2-GFP plasmid (provided by Prof. Philip Leder, Harvard Medical School). After purification using RNeasy MinElute Cleanup Kit (Qiagen), 300 pg of mFmn2-GFP capped RNA was co-injected with MO SB Fmn2 morpholino per embryo.

### Validation of splice blocking morpholino

The spice blocking morpholino MO SB Fmn2 was validated by RT-PCR. Total RNA isolated from morpholino injected 48 hpf embryos (RNeasy mini kit, Qiagen) was reverse transcribed using the SuperScript IV RT Kit (ThermoFisher). The cDNA obtained was amplified by the following primers flanking the intron between exon 5 and 6.

Primer sequences:

MOSBFmn2_RT_FWD: 5’-TCTGTTTGCATTGGGAGC-3’

MOSBFmn2_RT_REV: 5’-CTTGGTCTTTGACCTGCTGAT-3’

In control morphants, the expected amplicon size is 251 bp whereas in Fmn2 morphants the amplicon size was expected and found to be 550 bp due to blocked splicing of the intron between exons 5 and 6.

### Whole mount immunostaining, FM 4-64 labelling and fluorescence microscopy

PTU treated embryos were collected at desired stages and fixed in 4% PFA overnight at 4°C. The fixed embryos were stained as described earlier (Hatta, 1992) using 3A10 (DSHB) (1:50) antibody in blocking buffer. The larvae were washed with PBS-triton (0.5%) followed by blocking in 5% BSA. The larvae were then stained with anti-mouse AlexaFluor-568 (1:200) overnight at 4°C in blocking buffer. After extensive washing with PBS-triton (0.5%), the embryos were cleared in 50% glycerol and mounted dorsal side down on a glass bottom petri dish using low gelling agarose (Sigma).

To label the hair cells of the inner ear cristae, 1 nl solution of 3 μM FM 4-64 (Molecular Probes, Invitrogen) dissolved in DMSO was injected in the otic cavity of 96 hpf zebrafish embryos mounted laterally in low gelling agarose (Sigma). The injected embryos were removed from the gel using E3 buffer and imaged within 1 hour of injections. The samples were imaged on the LSM 780 confocal microscope (Zeiss) using a 25x oil immersion objective (NA 1.4).

### Behaviour experiment set up and behaviour analysis

We designed a behavioural assay as described previously (Lacoste et al., 2014) to screen larvae based on their response to a mechano-acoustic stimulus. The control and Fmn2 morphants were anaesthetized using 0.03% tricaine (MS-222, Sigma), head-restrained using 1% low gelling agarose and their tails were suspended in E3 buffer, in a 35 mm petri dish. All the fish were habituated for 30 minutes in the behaviour room maintained at a temperature of 27 °C. Individual dishes containing one larva were placed and taped onto the behavioural setup, as shown in Figure 2 A. Up to 6 taps were delivered using a 14 V DC solenoid (obtained from a local store) to the dish at an interval of 10 seconds at an intensity corresponding to 14 V from a power supply. The setup comprises of an automated stimulus delivery control unit (Arduino), a solenoid with a piston driven by a variable power supply, a piezo sensor (SparkFun) to detect the stimulus, and a feedback TTL pulse to the camera to mark the reception of the stimulus. This allowed precise marking of the stimulus delivery directly onto the images acquired using a high-speed video camera (AVT Pike, F-032B) at 640 fps. The secure image signature (SIS) feature of the camera was used to put a time-stamp on individual frames acquired. The recordings were analyzed using a custom-written Python program to extract the time-stamp, time of stimulus delivery and skeletonizing the fish to obtain coordinates of a spline curve fit to the fish shape in every frame. The skeleton was segmented into 20 points, and the last five points were used for further calculations. The program calculates the tail angle from last five points on the fish skeleton with respect to the restrained head segment. An event qualifies as an escape response if the angle crosses a threshold of 60 degrees from the rest position (Lacoste et al., 2014).

**Figure 1.**
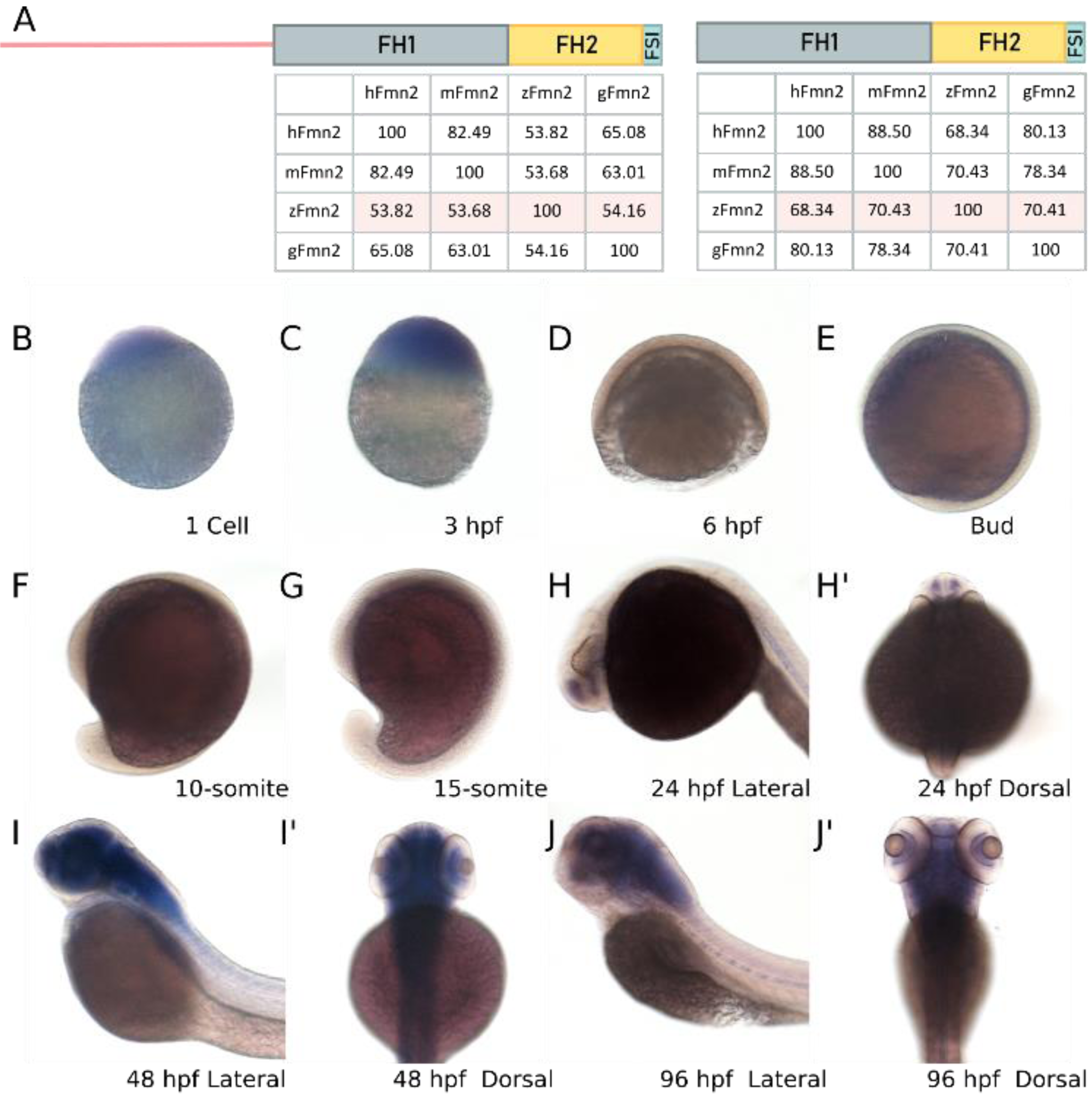
Fmn2 is maternally deposited and enriched in the zebrafish nervous system. Amino acid sequence alignments were obtained using EMBL Clustal Omega for human Fmn2 (hFmn2), mouse Fmn2 (mFmn2), zebrafish Fmn2 (zFmn2) and chicken Fmn2 (gFmn2) to compare conservation of sequence across vertebrate species. Percent identity matrices are shown for comparison of full length Fmn2 and the functional domains of Fmn2 across species. Full length zFmn2 showed 53.82%, 53.68% and 54.16% sequence similarity for human, mouse and chicken Fmn2 respectively. On comparing the functional domains of Fmn2, namely Formin Homology 1 (FH1), Formin Homology 2 (FH2) and Formin Spire Interaction domain (FSI), we found 68.34%, 70.43% and 70.41% sequence similarity with human, mouse and chicken Fmn2, respectively. **B-J)** Representative images of whole mount mRNA *in situ* hybridization showing the Fmn2 mRNA expression pattern in the developing zebrafish embryo. Fmn2 mRNA can be found in **B)** 1 cell stage and **C)** 3 hpf embryos suggesting maternal deposition. The mRNA expression is negligible during **D)** 6 hpf, **E)** bud stage, **F)** 10-somite and **G)** 15 somite embryos. Fmn2 mRNA expression is regained in **H)** 24 hpf embryos and continues to be expressed until 96 hpf **(I-J’)**.

**Figure 2.**
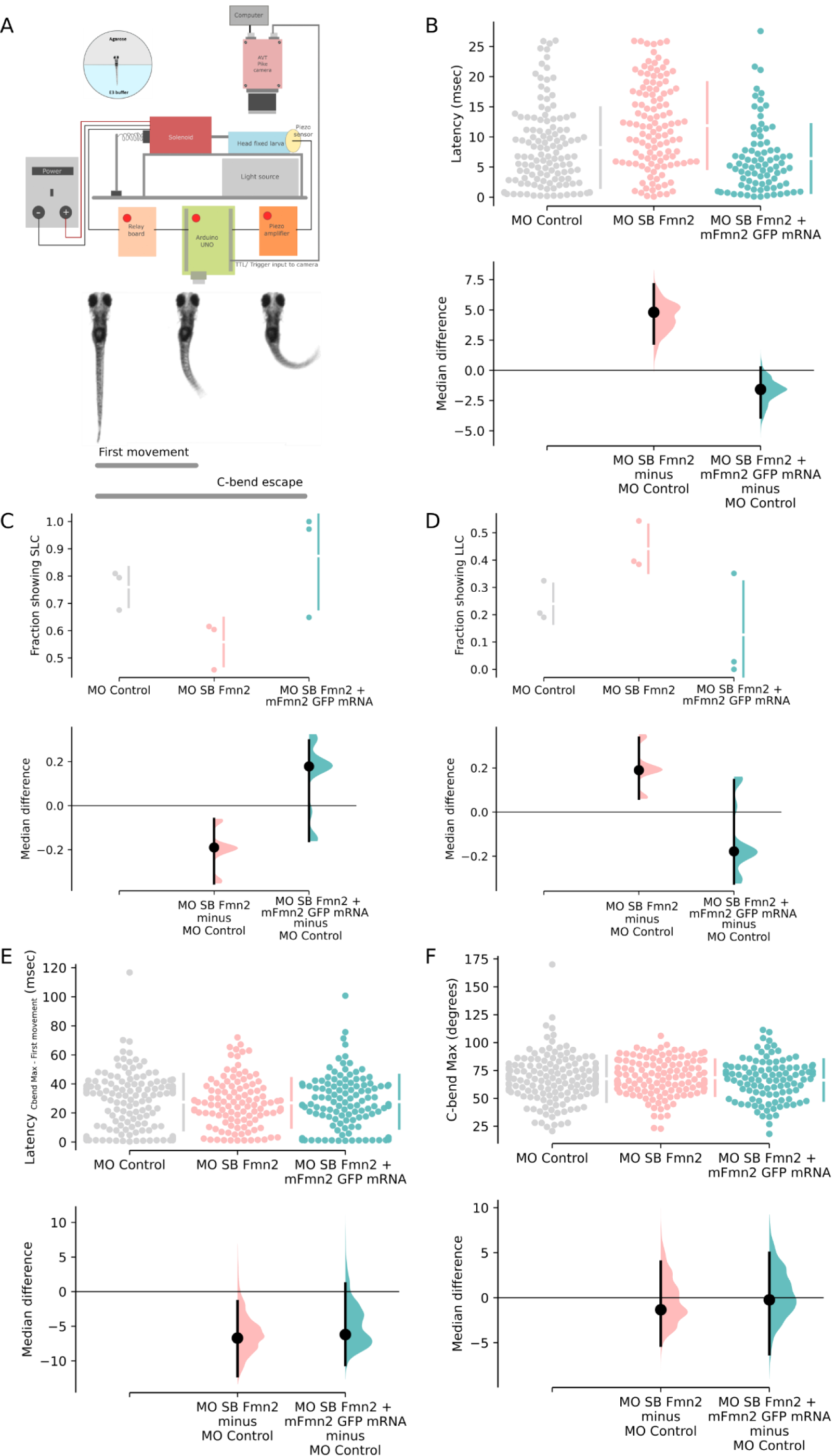
Fmn2 knockdown increased the latency to respond to mechano-acoustic stimulus. Schematic of the automated stimulus delivery apparatus and description of movements qualifying as the first movement for latency calculation and maximum C-bend for the escape response following the mechano-acoustic stimulus. **B-F)** The median difference comparisons against the control morpholino (MO Control) are shown in the Cumming estimation plots. The raw data is plotted on the upper axes as individual dots. The vertical gapped lines summarise the median ± standard deviation for each group. On the lower axes, median differences are plotted as bootstrap sampling distributions obtained by resampling the data 5000 times. The median difference is depicted as a dot for respective groups as compared to the shared control. The 95% confidence interval is indicated by the ends of the vertical error bars. Latency to first movement (msec) is plotted in this graph. The unpaired median difference between MO Control (n=126) and MO SB Fmn2 (n=120) is 4.8 [95.0%CI 2.23, 7.1]. The *P* value of the two-sided permutation t-test on the median differences is 0.0. The unpaired median difference between MO Control and MO SB Fmn2 + mFmn2-GFP mRNA (n=83) is -1.58 [95.0%CI -3.89, 0.232]. The *P* value of the two-sided permutation t-test on the median differences is 0.081. Values calculated for the fraction of the population (n=3) showing Short Latency C bend response (SLC) (< 13 msec) are plotted in this graph. The unpaired median difference between MO Control and MO SB Fmn2 is -0.201 [95.0%CI -0.308, -0.107]. The *P* value of the two-sided permutation t-test is 0.0. The unpaired median difference between MO Control and MO SB Fmn2 + mFmn2-GFP mRNA is 0.114 [95.0%CI - 0.151, 0.266]. The *P* value of the two-sided permutation t-test is 0.357. Values calculated for the fraction of the population (n=3) showing Long Latency C bend response (LLC) (13-26 msec) are plotted in this graph. The unpaired median difference between MO Control and MO SB Fmn2 is 0.201 [95.0%CI 0.104, 0.303]. The *P* value of the two-sided permutation t-test is 0.0. The unpaired median difference between MO Control and MO SB Fmn2 + mFmn2-GFP mRNA is -0.114 [95.0%CI - 0.276, 0.111]. The *P* value of the two-sided permutation t-test is 0.29. The difference of time taken to achieve the maximum C-bend angle and the latency to first movement for each trial is plotted in this graph. The unpaired median difference between MO Control and MO SB Fmn2 is -6.71 [95.0%CI -12.2, -1.4]. The *P* value of the two-sided permutation t-test is 0.0526. The unpaired median difference between MO Control and Rescue is -6.2 [95.0%CI -10.6, 1.17]. The *P* value of the two-sided permutation t-test is 0.152. This graph shows the maximum angle (C-bend max) attained during the escape response. The unpaired median difference between MO Control and MO SB Fmn2 is -1.34 [95.0%CI -5.31, 4.0]. The *P* value of the two-sided permutation t-test is 0.694. The unpaired median difference between MO Control and Rescue is -0.235 [95.0%CI -6.26, 4.98]. The *P* value of the two-sided permutation t-test is 0.875. For all the estimation statistics analysis, the effect sizes and CIs are reported as effect size [CI width lower bound; upper bound].

The following parameters have been quantified, as mentioned below:

#### Latency to first movement

Time taken by the fish to initiate movement post stimulus delivery, marked by an angle change greater than five degrees.

#### C-bend Max

Maximum angle of C-bend escape (with respect to the head restrained segment)

#### Latency to C-bend Max

Time taken by the fish to reach the maximum C-bend escape angle, calculated by subtracting latency to first movement from the total time taken to reach maximum angle.

### Figures and Statistical analysis

The images were processed in Fiji and assembled as figure panels using Inkscape. The data for all measurements are represented as (median; [95% confidence intervals]; number of events) for Figure 2 in the results section. Data obtained from various experiments were analyzed using an estimation statistics approach (Ho et al., 2019) and the median difference values and respective permutation test p-values are indicated in the figure legends. All data points have been presented as a swarm plot for individual values displaying the underlying distribution. The effect size is presented as a bootstrap 95% confidence interval (95% CI) below the swarm plots showing the median differences obtained by resampling the data 5000 times. The plots were generated using the web application available at https://www.estimationstats.com/#/.

## RESULTS

### The zebrafish ortholog of Fmn2 is enriched in the nervous system

Homology search comparing zebrafish sequences with human, mouse and chick Fmn2 cDNA sequences revealed a potential ortholog in zebrafish on Chromosome 12. We aligned the amino acid sequences of the orthologs using EMBL Clustal Omega. High sequence similarity was observed between the human, mouse, chick and zebrafish Fmn2 orthologs with the C-terminal FH1, FH2 and FSI domains showing ∽70% similarity (Fig. 1 A). Using whole mount *in situ* hybridization, we evaluated the expression pattern of Fmn2 mRNA in zebrafish across developmental stages (Fig. 1 B-J). We found that Fmn2 mRNA is maternally deposited, as seen in 1 cell stage zebrafish embryos, and the expression persists till 3 hours post fertilization (hpf). There was no discernible Fmn2 mRNA expression at stages from 6 hpf to 18 hpf. The mRNA expression reappeared in the telencephalon/forebrain at 24 hpf (Fig. 1 H). At stages 48 hpf onwards, the expression extends to the diencephalon/midbrain, the rhombencephalon/hindbrain, the spinal cord and the retinal ganglion cells (RGC) layer of the eye. Similar expression pattern is observed at 96 hpf (Fig. 1 I-J).

In line with previous studies showing the expression of Fmn2 in the nervous system of human, mice (Leader and Leder, 2000), and chick (Sahasrabudhe et al., 2016), Fmn2 mRNA is also enriched in the zebrafish nervous system.

### Fmn2 morphants exhibit a delay in initiating the escape response

To test the contribution of Fmn2 in development of the nervous system, we used a morpholino-mediated knockdown approach. We designed two Fmn2-specific antisense morpholinos, one translation blocking (MO TB Fmn2) and one splice blocking (MO SB Fmn2). The MO TB Fmn2 morpholino targets the first exon, and the MO SB Fmn2 blocks splicing at the boundary of the fifth exon and intron (Fig. 2-1 A). To test the extent of knockdown by MO SB Fmn2, RT-PCR was performed to ensure incorporation of the intron after exon 5, causing inclusion of a stop codon in the translation frame. In 48 hpf MO SB Fmn2 morphants, the majority of the Fmn2 mRNA was present in the splice-blocked form evident from the 550 bp amplicon corresponding to the inclusion of the intron between exons 5 and 6, as compared to the spliced version with an expected amplicon size of 251 bp (Fig. 2-1 B).

Out of all the embryos injected at 1 cell stage with MO TB Fmn2 and MO SB Fmn2, around 40% embryos exhibited morphological defects, like microcephaly, cardiac edema and axial curvature. These were excluded from further experiments. The larvae with intact otolith, inflated swim bladders and length comparable to the control morphants were used for all analyses.

To evaluate the effect of Fmn2 knockdown on behaviour, we recorded the responses of control and Fmn2 morphants to manual tapping of the dish containing the larvae. In this preliminary experiment, we observed uncoordinated locomotion in Fmn2 morphants. To better resolve these behavioural defects, we employed high speed video recording and tested the response of morphants to mechano-acoustic stimuli. In this assay, we subjected 96 hpf head restrained morphants to a tap on the dish in which they were housed and measured their response by tracking the movement of their tails. The response of the morphants was recorded at 640 fps using a high-speed video recording camera (AVT Pike, F-032B). The taps were delivered at 10 second intervals, controlled by an Arduino UNO microprocessor (Fig. 2 A). The recorded videos were analyzed using a custom written python program to extract time-stamp, and the body axis coordinates after skeletonizing the shape of the fish (Movie 1).

The latency to first movement, the maximum escape angle (C-bend Max) and the latency to achieve maximum angle in an escape response (Latency _C-bend Max - First movement_) were calculated.

The tap strength using the piston was controlled by varying the input voltage on the variable power supply. For all the experiments, tap strength corresponding to 13 volts was used. This tap strength was optimized to achieve about 95% responsiveness in the control zebrafish larvae. The responsiveness of the Fmn2 morphants (injected with 2 ng/embryo MO SB Fmn2) to the same mechano-acoustic stimulus was comparable (94%) to the control animals (Fig. 2-1 C). On the other hand, we observed that Fmn2 morphants showed an increased latency to first movement (11.54 msec; [9.36,13.38]; n=120; Fig. 2 B) in response to the mechano-acoustic stimulus as compared to the control morphants (6.74 msec; [5.39,9.06]; n=126; Fig. 2 B). To ensure that the morpholino’s effect was specific to Fmn2, we co-injected mouse Fmn2-GFP (mFmn2-GFP) mRNA, which is resistant to the anti-zebrafish Fmn2 morpholino, along with the Fmn2 morpholinos. The increased latency could be rescued in Fmn2 morphant larvae co-injected with mFmn2-GFP mRNA (5.16 msec; [4.03,6.29]; n=83; Fig. 2 B). Similar defects were observed in Fmn2 morphants injected with MO TB Fmn2 (17.7 msec; [16.50,19.00]; n=82; Fig. 2-1 C).

Previous reports implicate Mauthner cells (M-cell) in mediating short latency (SLC; <13 msec) escapes in response to mechano-acoustic stimuli (Lacoste et al., 2014). Whereas, long latency escapes are generally non-Mauthner mediated responses. We classified the latency to the first movement as short latency C-bend (SLC; <13 msec) and long latency C-bend events (LLC; 13-26 msec). Of the total number of events, we observed a decrease in the fraction of SLC versus LLC responses in Fmn2 morphants.

In morphants co-injected with mFmn2-GFP mRNA, the fraction of events showing SLC responses is greater than LLC responses as seen in control morphants (Fig. 2 C, D). However, the latency to achieve maximum C-bend escape angle (Fig. 2 E) and the maximum escape angle (Fig. 2 F) were comparable between the control morphants, the Fmn2 morphants and the Fmn2 morphants co-injected with mFmn2 mRNA.

Hence, Fmn2 knockdown increased the latency to respond to mechano-acoustic stimuli even though the responsiveness and the ability to elicit an escape response remains uncompromised. These results suggest a specific and perhaps localized defect in the mechano-acoustic response circuitry.

### Fmn2 depletion does not affect the sensory components of the acoustic startle circuit

The acoustic startle response is essential for the survival of zebrafish larvae in terms of reacting to the environment efficiently and reliably. The behavioural deficits observed in Fmn2 morphants could arise from one or more components of the acoustic startle circuit. We systematically evaluated the different components of the acoustic startle circuit to identify the origins of the delay in the initiation of the C-bend escape in Fmn2 morphants. M-cells receive auditory input from the inner ear hair cells, relayed by the statoacoustic ganglion (SAG) (Whitfield et al., 2002; Medan and Preuss, 2014). Evaluation of the otic vesicle and the otoliths did not reveal any anatomical or structural defects. We probed the structural integrity of the inner ear hair cells responsible for the mechanotransduction of the acoustic cues, using the Tg(Brn3c:GAP43-GFP) line. This line labels a subset of the hair cells in the inner ear and lateral line neuromasts (Xiao et al., 2005). Microscopic analysis of kinocilia and hair cell bundles of the inner ear cristae revealed no significant differences between control and Fmn2 morphants (Fig. 3 A-D). Uptake of the lipophilic dye FM 4-64 was used to evaluate the functional activity of the inner ear hair cells (Pacentine and Nicolson, 2019). These experiments indicated that activity-dependent vesicle recycling was comparable between control and Fmn2 morphants (Fig. 3 E, F) and suggested that the synaptic activity at the hair cell ribbon synapses is mostly unaffected in the Fmn2 morphants.

**Figure 3.**
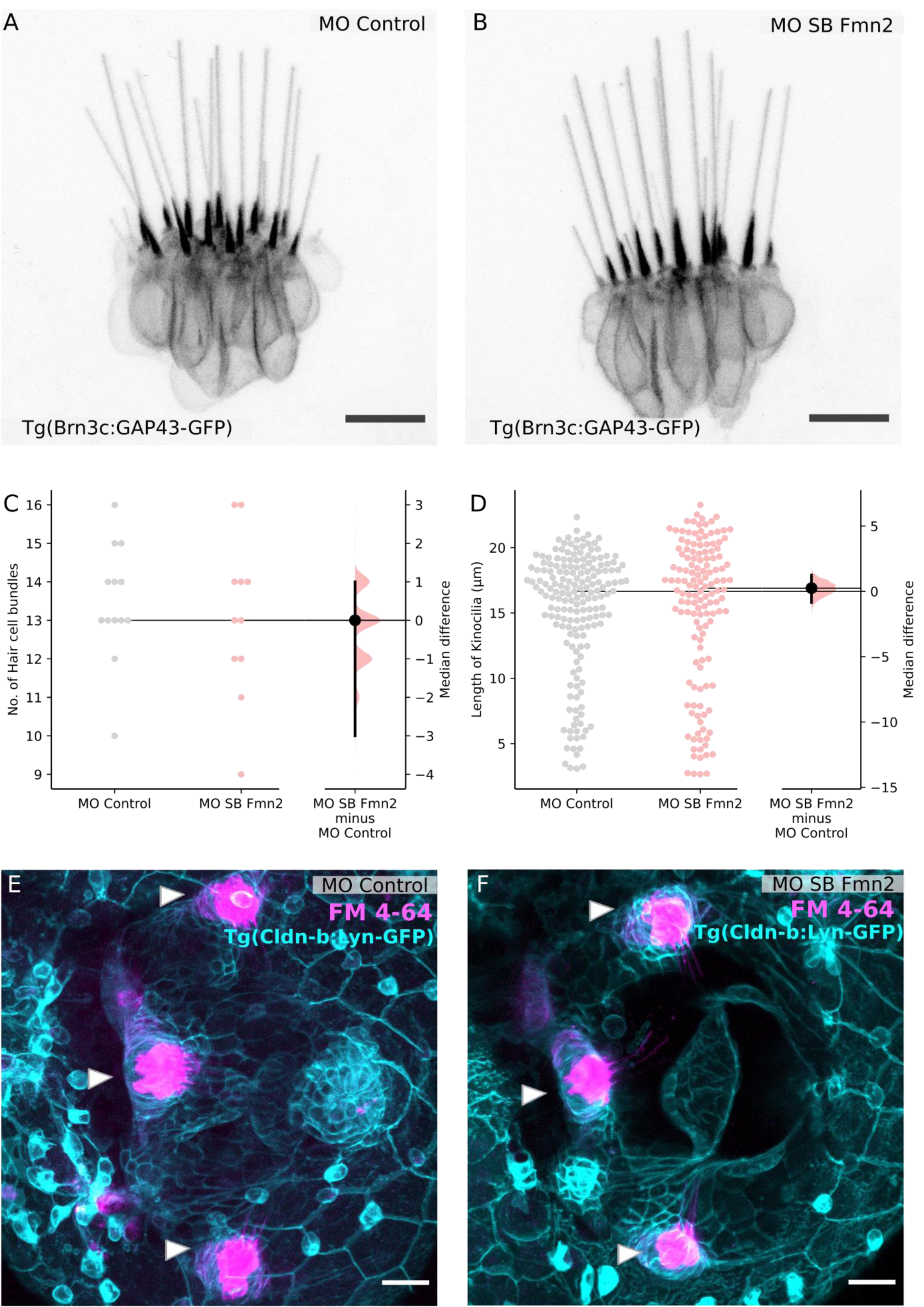
Sensory components of the acoustic startle circuit are not affected in Fmn2 morphants. Hair cells of the lateral crista of the zebrafish inner ear are visualized using the Tg (Brn3c:GAP43-GFP) line in 96 hpf larvae with **A)** 2 ng MO Control and **B)** 2 ng MO SB Fmn2 cytoplasmic injections. Scale bar is equivalent to 10 μm. Quantification of the number of hair cell bundles with a kinocilium is depicted in the Gardner-Altman plot. The unpaired median difference between MO Control and MO SB Fmn2 is 0.0 [95.0%CI -3.0, 1.0]. The P value of the two-sided permutation t-test is 0.687. Quantification of kinocilia length in the lateral crista hair cells is shown in the Gardner-Altman plot. The unpaired median difference between MO Control and MO SB Fmn2 is 0.247 [95.0%CI -0.851, 1.25]. The *P* value of the two-sided permutation t-test is 0.663. For all the estimation statistics analysis, the effect sizes and CIs are reported as effect size [CI width lower bound; upper bound]. Representative images for FM-4-64 dye uptake assay in the inner ear of 96 hpf **E)** control morphants and **F)** Fmn2 morphants, done in the background of Tg(Cldn-b:lyn-GFP) to mark the inner ear boundary. Scale bar is equivalent to 20 μm.

The next component of the acoustic startle circuit is the Statoacoustic ganglion (SAG) which connects the inner ear hair cells to the M-cell. SAG was also found to be structurally unperturbed as evident in 96 hpf morphants immunostained with anti-neurofilament 3A10 antibody (Fig. 4 A, B).

**Figure 4.**
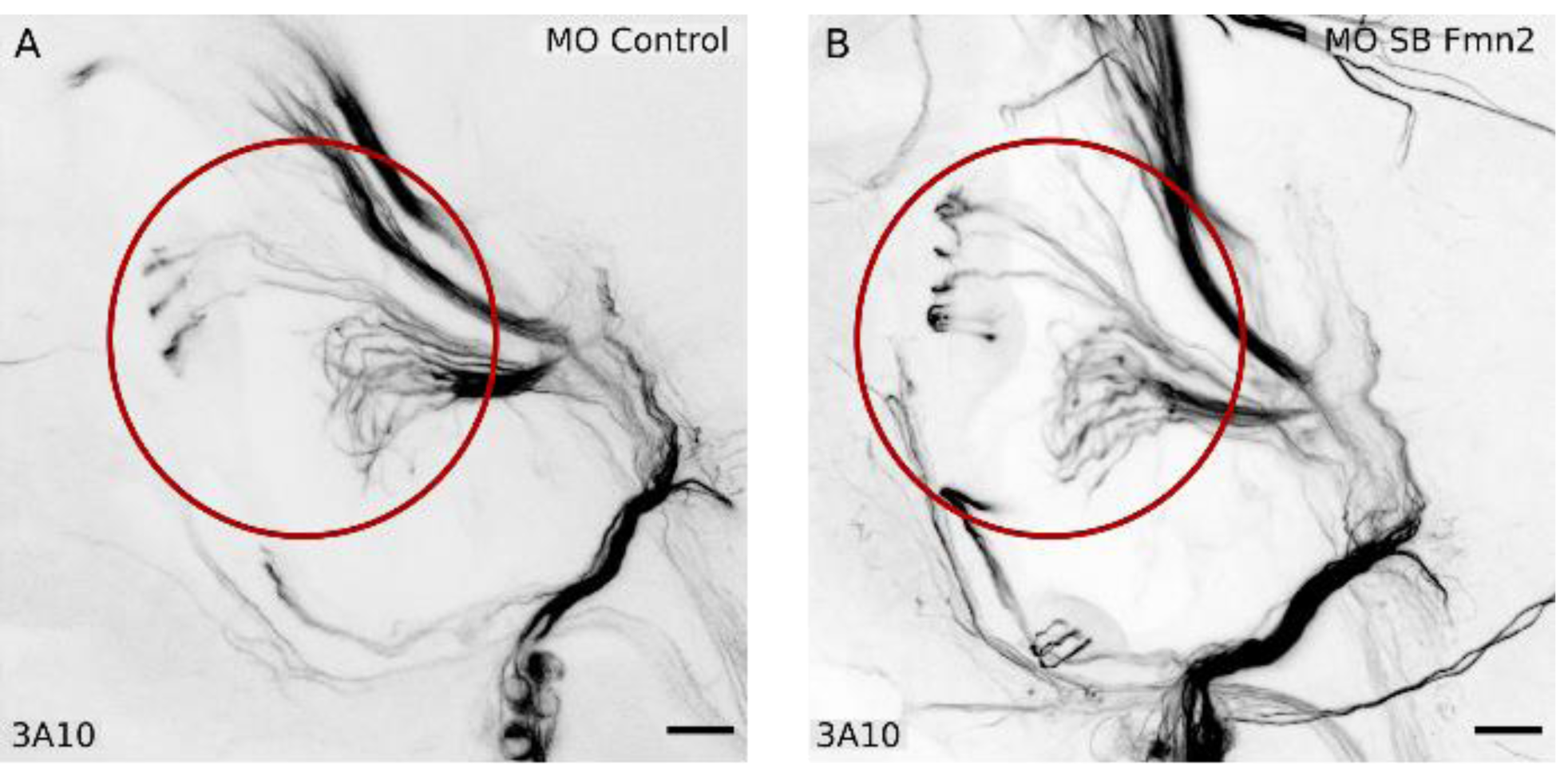
The Statoacoustic ganglion (SAG) relaying sensory information is not affected in Fmn2 morphants. Whole mount immunostaining with anti-neurofilament antibody 3A10 was used to visualize the Statoacoustic ganglion connecting to the M-cells in 96 hpf larvae with **A)** 2 ng MO Control and **B)** 2 ng MO SB Fmn2 cytoplasmic injections. Scale bar is equivalent to 20 μm.

Our findings suggest that the sensory components of the acoustic startle circuit providing input to the M-cell are unaffected by Fmn2 knockdown. This observation is consistent with the unaffected responsiveness to mechano-acoustic stimuli in Fmn2 morphants.

These results suggest that the behavioural defect of increased latency is caused by deficits in the neural circuit downstream of the statoacoustic ganglion. This encouraged us to probe the locomotion circuitry in the zebrafish hindbrain.

### Spiral fiber neuron development in the hindbrain is regulated by Fmn2

The M-cell is known to receive inputs from excitatory as well as inhibitory interneurons (Korn and Faber, 2005) to achieve modular fine-tuning of responses to a variety of stimuli. The increase in the response latency in Fmn2 morphants suggests possible defects in the hindbrain circuits involving the M-cells. To assess the state of neuronal connectivity in the hindbrain, we visualized the axonal tracts in 96 hpf morphants using the anti-neurofilament antibody 3A10. We found that the majority of the axonal tracts in the hindbrain, including the M-cell and its homologs, remain unaffected in Fmn2 morphants. However, the axonal tracts of the Spiral fiber neurons in rhombomere 3 were absent. Spiral fiber neurons are commissural interneurons innervating the M-cell axon hillock (Scott et al., 1994; Lorent et al., 2001; Gyda et al., 2012; Lacoste et al., 2014; Liu and Hale, 2017). They have been described earlier to provide excitatory feedforward input to the M-cells and regulate the escape response in larval zebrafish (Lacoste et al., 2014). We found that 48% of the MO SB Fmn2 morphants (n=31) showed absence of spiral neurons, whereas none of the control morphants (n=27) showed the defect (Fig. 5 D). The phenotype persisted with the second morpholino, MO TB Fmn2 (Fig. 5-1 A). We were also able to rescue the phenotype by co-injecting mFmn2-GFP mRNA along with MO SB Fmn2. The phenotype was rescued in 95% (n=40) of the morpholino and mFmn2-GFP mRNA co-injected embryos.

**Figure 5.**
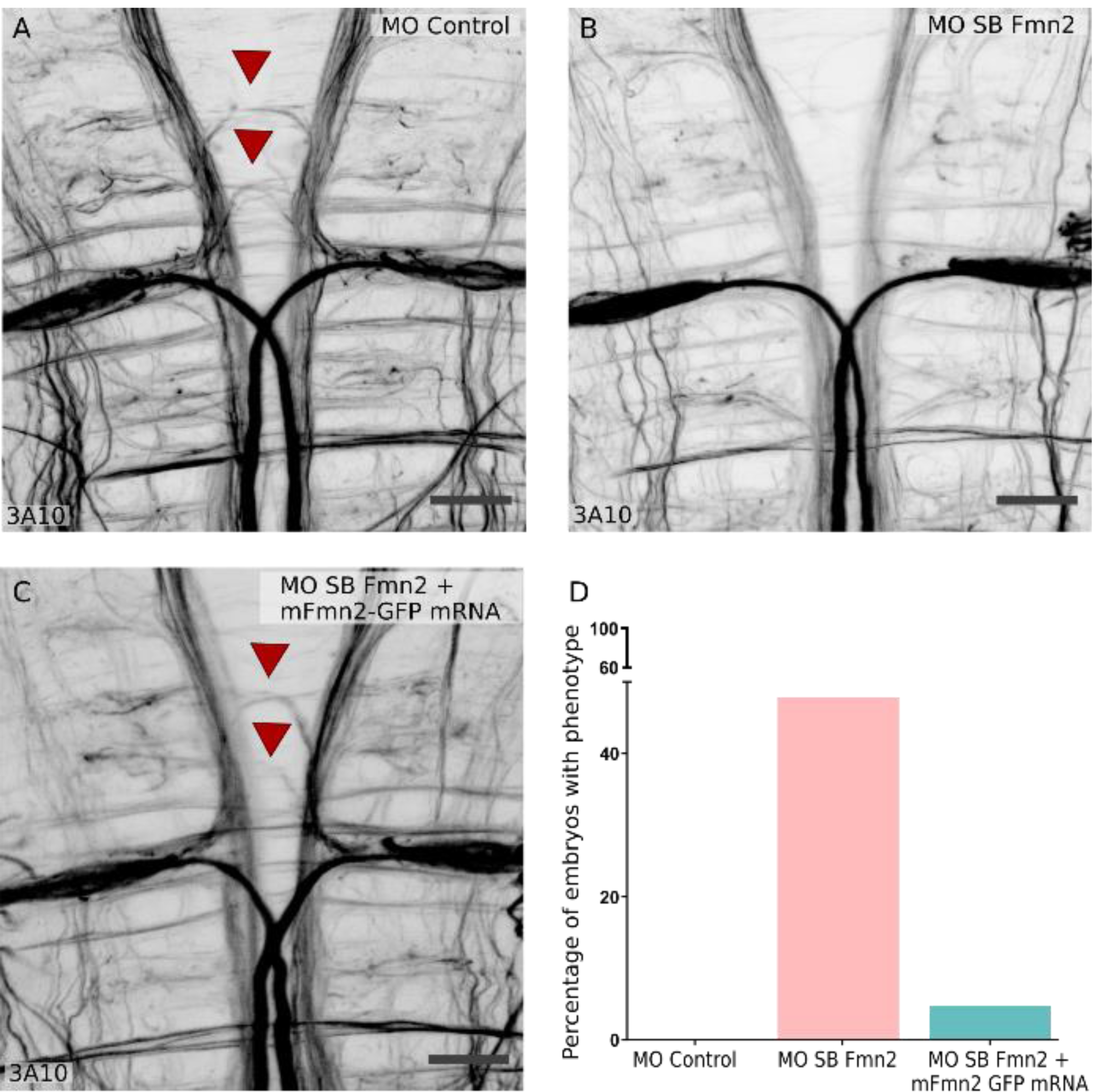
Fmn2 knockdown impairs axonal outgrowth in the spinal fiber neurons. Whole mount immunostaining using anti-neurofilament antibody 3A10 stains axons in 96 hpf zebrafish hindbrain. **A)** Spiral fiber neurons are marked with red arrowheads in control morphants. B) Fmn2 knockdown using splice blocking morpholino (MO SB Fmn2; 2 ng/embryo injected in the cytoplasm) reveals defects in the spiral fiber neurons outgrowth. C) The phenotype in Fmn2 morphants could be rescued by injection of 300 pg mFmn2-GFP mRNA in the MO SB Fmn2 injected embryos at 1 cell stage. Scale bar is equivalent to 20 μm. D) A total of 48% Fmn2 morphants (n=31) show the absence of spiral fiber neurons as compared to none in the control morphants (n=27). Whereas, only 5% of the embryos rescued with mFmn2-GFP mRNA (n=40) showed the defects.

These results implicate Fmn2 in the development of the spiral fiber neurons that are necessary for the efficient and reliable execution of the M-cell mediated C-bend escape response. The increase in latency to respond to mechano-acoustic stimulus in Fmn2 morphants can be attributed to the lack of innervation of the M-cells by excitatory spiral fiber neurons.

## DISCUSSION

The zebrafish Fmn2 ortholog was identified to be located on chromosome 12. The three key functional protein domains at the C-terminus (FH1, FH2 and FSI domains) that define the fly, chick, mouse and humans orthologs of Fmn2 (Higgs, 2005; Breitsprecher and Goode, 2013) were found to be highly conserved in zebrafish. On the other hand, the N-terminal region was less conserved across these species, perhaps reflecting species-specific regulatory diversity. However, the ability of mouse Fmn2 mRNA to rescue the Fmn2 depletion induced phenotypes in zebrafish underscores the functional conservation between these species.

Using reciprocal BLASTp, we found that E7F517 (UNIPROT) has the highest sequence identity with human Fmn2 (53.82 % Fig. 1 A). The next best hit, X1WC43 (UNIPROT), was found to have considerably less sequence identity with human Fmn2 (44.75 %). X1WC43 is located on chromosome 17 and has a characteristic FH2 domain, but the FH1 domain is truncated to 33 amino acids. Further, mRNA corresponding to X1WC43 was found to have no discernible expression in the zebrafish nervous system (data not shown). Therefore, we identify zebrafish E7F517 to be the functional ortholog of human Fmn2.

Fmn2 mRNA is enriched in the developing zebrafish nervous system though the expression pattern is dynamic during early development. Fmn2 mRNA is maternally deposited in embryos but disappears after 3 hpf. Robust expression of Fmn2 mRNA in the nervous system resumes by 24 hpf and persists till 96 hpf (Fig. 1 B-J’). The spatiotemporal expression pattern coincides with neuronal development in the zebrafish embryo.

Behavioural analysis using an automated stimulus delivery setup and high-speed recording revealed a specific function of Fmn2 in the acoustic startle response. The responsiveness of Fmn2 and control morphants were comparable and indicated that sensory perception was unaffected. However, Fmn2 knockdown increased the latency to respond; the proportion of fast responses decreased while the long latency escape responses increased (Fig. 2 B-D).

Consistent with the behavioural observations, the inner ear hair cells (Fig. 3 A-F) and the statoacoustic ganglion (Fig. 4 A-B) were found to be unaffected by Fmn2 depletion. Strikingly, we found that spiral fiber neurons, which provide excitatory input to the M-cells, fail to form synaptic terminals at the M-cell axon hillock (Fig. 5 A-B). Thus the observed deficits in behaviour are likely due to the absence of spiral fiber neuron innervation resulting in the failure of the command neuron-like M-cells to reach the excitatory threshold (Lacoste et al., 2014). We were able to rescue both the behavioural (Fig. 2 B-D) and the neuro-anatomical phenotypes in Fmn2 morphants by co-injection of mFmn2-GFP mRNA (Fig. 5 C). This result underscores the evolutionarily conserved function of Fmn2 from teleosts to mammals.

Previous studies have highlighted the role of spiral fiber neurons in regulating the fast escape responses in response to mechano-acoustic stimuli (Lorent et al., 2001; Lacoste et al., 2014; Hale et al., 2016; Marsden et al., 2018). Spiral fiber neurons directly respond to mechano-acoustic stimuli and relay the information to the M-cells, which also receive direct sensory inputs (Lacoste et al., 2014). In a separate study, the absence of spiral fiber neurons along with other hindbrain commissures caused locomotor defects in *space cadet* mutants (Lorent et al., 2001) later characterized as a mutation in the retinoblastoma-1 (Rb1) gene (Gyda et al., 2012). These studies indicate that the spiral fiber neurons are indispensable for the initiation of a robust startle escape response (Hale et al., 2016). The convergent circuit design in zebrafish hindbrain, where spiral fiber neurons help M-cells reach the excitatory threshold, ensures a speedy and reliable response to a potentially noxious stimulus. The excitability of M-cells decides the further course of action for the animal within the specific context.

Zebrafish need the M-cells and two segmental homologs, MiD2cm and MiD3cm, to elicit an effective, fast escape response in zebrafish (Liu and Fetcho, 1999; Kohashi and Oda, 2008). These three neurons receive a common auditory input but can generate different outputs, evident by their different spiking properties, to downstream neurons to control adaptive escape behaviours. MiD2cm and MiD3cm mediate escapes with longer latencies as compared to the M-cell (Nakayama and Oda, 2004; Kohashi et al., 2012; Lacoste et al., 2014). The M-cell homologs ensure that an escape response is elicited in the absence of M-cell firing or for weaker stimuli. Ablation of the M-cell, especially the axon initial segment, causes an increase in latency of response to mechano-acoustic stimuli (Liu and Fetcho, 1999; Hecker et al., 2020).

In Fmn2 morphants, the M-cells fail to receive inputs from the spiral fiber neurons and exhibit a delay but eventually elicit an escape response. It is likely that MiD2cm and MiD3cm neurons, which also receive the auditory input from the statoacoustic ganglion, are recruited in Fmn2 morphants to produce an escape response with increased latency. In addition to the overall increase in latency in Fmn2 morphants, the shift towards majority LLC responses in Fmn2 morphants instead of SLC responses implies the involvement of M-cell homologs in the absence of spiral fiber excitation of the M-cell.

While the possibility of further deficits downstream of the M-cell homologs cannot be ruled out, the Fmn2 morphants can still execute C-bend escapes with no significant changes in the maximum bending angle in response to mechano-acoustic stimuli. Therefore, we conclude that the absence of M-cell innervation by the spiral fiber neurons are the primary reason for the behavioural deficits in Fmn2 morphants.

Regulated and adaptive remodelling of the neuronal cytoskeleton is central to almost all aspects of neural development, including neurogenesis, neurite initiation, growth cone-mediated pathfinding and synaptogenesis (Lowery and Van Vactor, 2009; Flynn, 2013; Gordon-Weeks and Fournier, 2014; Muñoz-Lasso et al., 2020). The dynamics of the neuronal cytoskeleton is mediated by a complex interplay between the different cytoskeleton components, each regulated by specific regulators but also coordinated by co-regulatory activities. The dynamics actin cytoskeleton is regulated by several actin-binding proteins, and mutations in many of these have been associated with neurodevelopmental disorders (Lian and Sheen, 2015; Muñoz-Lasso et al., 2020).

Actin nucleators are an important class of actin-binding proteins that are involved in regulating the seeding of F-actin filaments from monomeric actin and control the architecture of the F-actin network. Three classes of actin nucleators have been described previously in zebrafish, Arp2/3 complex, Formin homology domain 2 (FH2) containing family (the Formins) and the WASP homology domain 2 (WH2) containing family of nucleators. Arp2/3 has been shown to regulate actin patches essential for proximal axon specification in zebrafish motor neurons (Balasanyan et al., 2017) and maintenance of microridge structure and length on surface epithelial cells in zebrafish (Lam et al., 2015; Pinto et al., 2019). The WH2 domain containing actin nucleator, Cordon bleu (Cobl), is required for the development of motile cilia in zebrafish Kuppfer’s vesicles which in turn maintain laterality in zebrafish (Ravanelli and Klingensmith, 2011).

There are fifteen formins in humans that cluster into eight different subfamilies (Schönichen and Geyer, 2010) and mutations in several formins are associated with a variety of neural disorders (Kawabata Galbraith and Kengaku, 2019). Most formins are conserved across vertebrates and are expressed in a variety of tissues, including the nervous system (Dutta and Maiti, 2015). The formin family comprises of 25 predicted members in zebrafish out of which 11 are shown to be neuronally enriched (Thisse and Thisse, 2005; Santos-Ledo et al., 2013). A study comparing expression patterns of the formin-like (fmnl) subfamily of formins shows that the formins, fmnl1a, fmnl1b, fmnl2a, fmnl2b and fmnl3 are expressed in the nervous system in a dynamic temporal manner during development, in addition to expression in non-neuronal tissues (Santos-Ledo et al., 2013). However, the role of formins in the development of neural circuits in zebrafish is not well explored despite several studies reporting the enrichment of formins in vertebrate nervous systems (Leader and Leder, 2000; Krainer et al., 2013; Dutta and Maiti, 2015; Sahasrabudhe et al., 2016). The only report available implicates the formin Daam1a in the asymmetric morphogenesis of the zebrafish habenula and the regulation of the dendritic and axonal processes of the dorsal habenular neurons (Colombo et al., 2013). Formin function in non-neuronal tissues has also been sparsely investigated in zebrafish. Daam1a is required for convergent extension and regulates notochord development in zebrafish (Kida et al., 2007) while zDia2, working synergistically with Profilin I, regulates gastrulation (Lai et al., 2008). Fmnl3 has been reported to be involved in blood vessel morphogenesis (Phng et al., 2015).

Formins are expressed in several tissue types with rich spatiotemporal diversity in humans (Krainer et al., 2013), mice (Dutta and Maiti, 2015) and zebrafish (Thisse and Thisse, 2005; Kida et al., 2007; Lai et al., 2008; Santos-Ledo et al., 2013; Sun et al., 2014). The conservation of multiple of formins with overlapping expression in the nervous system possibly reflects a diversity of distinct regulatory functions, ensuring the highly adaptive yet specific cytoskeleton remodelling necessary for accurate circuit development.

Given the broad expression of the Fmn2 mRNA in the developing and adult zebrafish nervous system, it is interesting to note that the defects due to Fmn2 knockdown are confined to a small population of hindbrain excitatory interneurons, the spiral fiber neurons. The axonal outgrowth of spiral fiber neurons is completed by 72 hpf (Lorent et al., 2001). The developmental timing of axonal outgrowth in this neuronal population and protein perdurance from the maternal Fmn2 mRNA may render spiral fiber neurons, especially sensitive to morpholino-mediated knockdown. There is a notable expression of Fmn2 mRNA in the retinal ganglion cells of the eye and the spinal cord of zebrafish larvae. However, the role of Fmn2 in development of other neural circuits in zebrafish remains untested.

Fmn2 has previously been shown to be necessary for the axonal outgrowth of spinal neurons in developing chick embryos (Sahasrabudhe et al., 2016; Ghate et al., 2020). Similarly, we speculate that the zebrafish spiral fiber neurons have axonal outgrowth defects that result in the loss of synaptic connectivity with the M-cells. The molecular mechanism mediating Fmn2-dependent axonal outgrowth in zebrafish is not known. In chick spinal neurons, Fmn2 mediates growth cone motility by regulating the cell-matrix adhesions necessary to generate traction forces (Sahasrabudhe et al., 2016; Ghate et al., 2020). A recent study implicates Fmn2 in regulating growth cone microtubule dynamics in zebrafish Rohon-Beard neurons (Kundu et al., 2020) and highlights another mechanism mediating outgrowth and pathfinding. However, it remains possible that there are additional deficits in neuronal differentiation or specification in the Fmn2 morphants. Further studies in zebrafish, involving *in vivo* imaging of cytoskeletal dynamics, can pave the way for mechanistic insights into Fmn2 function in intact animals.

In recent years, mutations in Fmn2 have been increasingly associated with neural disorders including cognitive disabilities and sensory processing dysfunction in humans (Perrone et al., 2012; Almuqbil et al., 2013; Law et al., 2014; Agís-Balboa et al., 2017; Anazi et al., 2017; Marco et al., 2018; Gorukmez et al., 2020). Fmn2 expression was found to be reduced in post mortem brain samples of patients with post-traumatic stress disorder and Alzheimer’s disease (Agís-Balboa et al., 2017). Other reports implicate Fmn2 mutations in corpus callosum agenesis (Perrone et al., 2012; Gorukmez et al., 2020) and microcephaly (Anazi et al., 2017). In rodents, loss of Fmn2 accelerated age-associated memory impairment and amyloid-induced deregulation of gene expression (Peleg et al., 2010; Agís-Balboa et al., 2017). Despite accumulating evidence, little is known about the function of Fmn2 in the nervous system. A zebrafish Fmn2 loss-of-function model can provide valuable neurodevelopmental insights into Fmn2 function.

This study explores Fmn2 function in a vertebrate model and links the axonal development of an identified group of neurons to a specific behavioural deficit. Fmn2 was found to mediate the development of the spiral fiber neuron pathway conveying indirect excitatory inputs to the command neuron-like M-cells. The loss of this specific regulatory unit is manifested in a delay in initiating fast escape reflexes in response to mechano-acoustic stimuli, a behaviour of significant survival value. Apart from identifying a novel function for Fmn2 in the development of hindbrain commissural circuitry, our findings highlight the utility of models bridging subcellular functions of Fmn2 identified in cultured neurons to circuit development and associated behavioural consequences.

## ETHICS APPROVAL

All protocols used in this study were approved by the Institutional Animal Ethics Committee and the Institutional Biosafety Committee of IISER Pune.

## AVAILABILITY OF DATA AND MATERIAL

All data generated or analyzed during this study are included in this published article. The raw data are available from the corresponding author on reasonable request.

## AUTHOR CONTRIBUTIONS

Conceptualization: D.N. and A.G.; Investigation and analysis: D.N. (all experiments and analysis), T.K.J. and D.N. (development of custom code for time-stamp extraction and behaviour analysis), R.M. (initial experiments with the translation blocking MO), S.G. (optimization of the *in situ* probe used in this study); Writing and editing of manuscript: D.N. and A.G.; Funding Acquisition: A.G.. All authors gave final approval for publication and agreed to be held accountable for the work performed therein.

## ACKNOWLEDGMENTS

The authors are grateful to Dr. Joby Joseph (Hyderabad Central University, India) for assisting in the development of the automated stimulus delivery apparatus and Dr. Chinmoy Patra (Agharkar Research Institute, Pune) for technical support on the whole mount *in situ* hybridization studies. Dr. Mahendra Sonawane (Tata Institute of Fundamental Research, Mumbai) is acknowledged for providing the Tg(Cldn-b:Lyn-GFP) line. The Tg(Brn3c:GAP43-GFP) line was a gift from Dr. Shawn Burgess (National Human Genome Research Institute, National Institutes of Health, USA). The authors thank Dr. N. K. Subhedar, IISER Pune and Dr. Raghav Rajan for critical inputs on the manuscript. The authors acknowledge the IISER Pune Microscopy Facility and the National Facility for Gene Function in Health and Disease (NFGFHD) at IISER Pune for access to equipment and infrastructure.

## FIGURES

**Movie 1. Tracking the escape response in a 96 hpf wildtype zebrafish larva**

The movie shows a wildtype larva being tracked by a custom written code which extracts time-stamp from the secure image signature on the frames and skeletonizes the fish to obtain coordinates for further calculations.

