## Supplemental Data for "The formin Fmn2 is required for the development of an excitatory interneuron module in the zebrafish acoustic startle circuit"

**This file includes:**

Figures 2-1 and 5-1

**Other supplementary materials for this manuscript include the following:**

Movie 1

### EXTENDED DATA

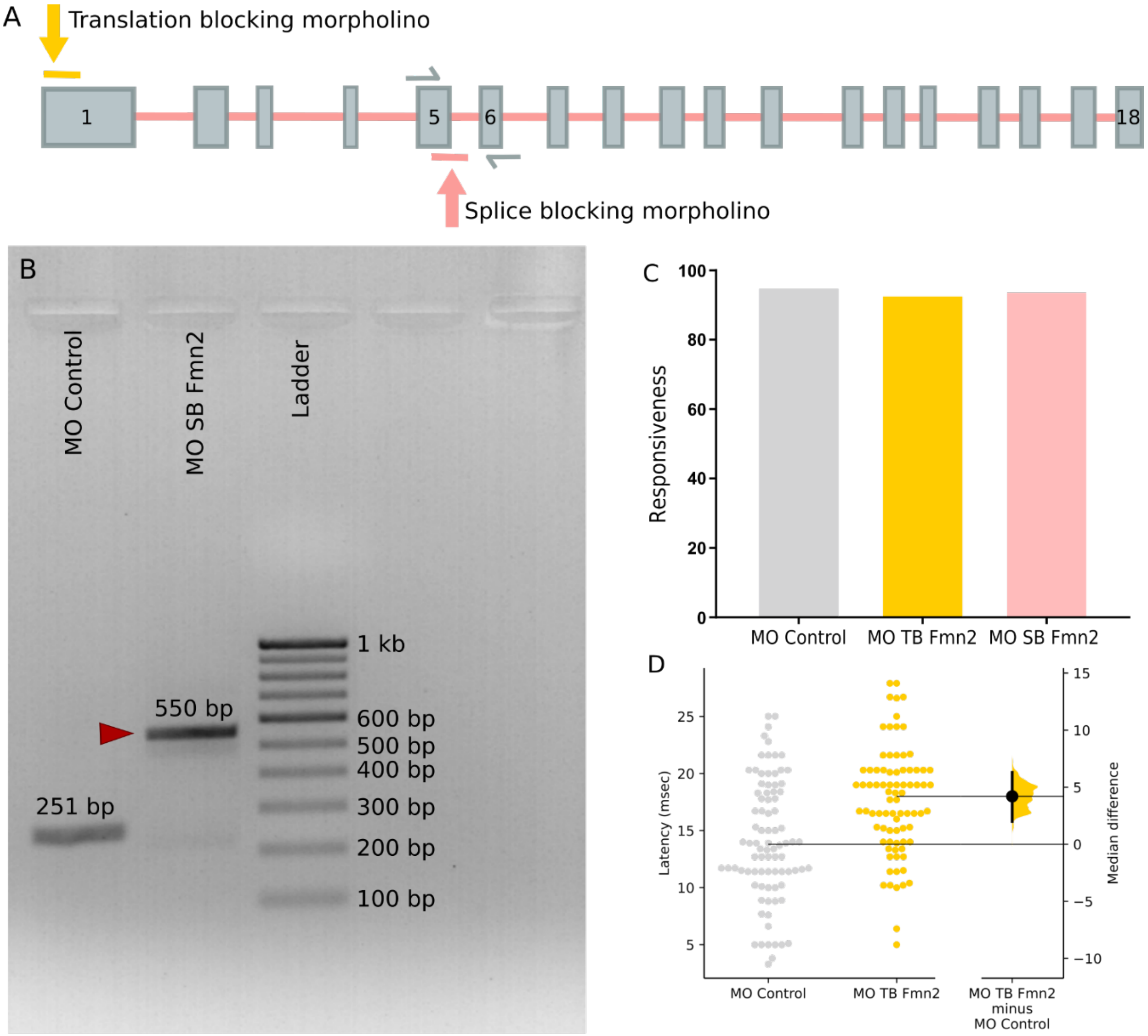


Extended Data Figure 2-1

**Figure 2-1. Morpholino design and validation**

**A)** Schematic showing target regions for the two morpholinos used in the experiments. MO TB Fmn2 targets exon 1 to block translation. MO SB Fmn2 targets the exon 5 – intron 5 boundary to cause retention of intron 5 between exons 5 and 6, leading to the occurrence of a premature stop codon. Both morpholinos ensure that the functional domains are not translated in Fmn2 morphants.

**B)** Validation of knockdown by MO SB Fmn2 morpholino was done using RT-PCR on cDNA obtained from MO Control and MO SB Fmn2 injected embryos. The amplification of a 550 bp amplicon from MO SB Fmn2 morphants cDNA corresponds to inclusion of intron 5 because of efficient splice blocking by the morpholino.

**C)** Responsiveness is quantified as the percentage of larvae responding to acoustic stimuli in MO Control (95.2 %), MO TB Fmn2 (92.9 %) and MO SB Fmn2 (94 %) injected embryos.

**D)** Behavioural analysis of MO TB Fmn2 morphants is summarized in the Cumming plot. MO TB Fmn2 morphants also exhibit increased latency defect. The unpaired median difference between MO Control and MO TB Fmn2 is 4.2 [95.0%CI 2.0, 6.3]. The *P* value of the two-sided permutation t-test is 0.0004. The effect sizes and CIs are reported as effect size [CI width lower bound; upper bound].

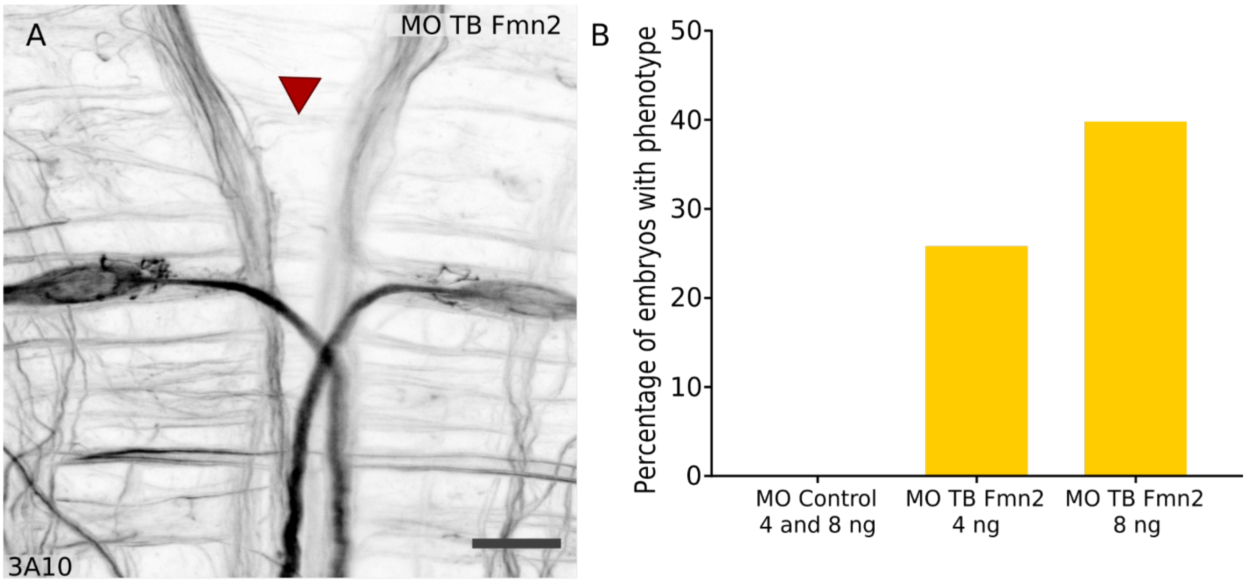


Extended Data Figure 5-1

**Figure 5-1. The translation blocking morpholino recapitulates the spiral fiber neuron defect in Fmn2 morphants**

**A)** Whole mount immunostaining using 3A10 antibody of 96 hpf Fmn2 morphants injected with 2 ng MO TB Fmn2 phenocopies the splice blocking Fmn2 morphant defects. Scale bar is equivalent to 20 μm.

**B)** In addition to cytoplasmic injections of MO TB Fmn2, yolk injections of higher doses (4 ng and 8 ng) of MO TB Fmn2 also cause the spiral fiber neuron outgrowth defect in a dose dependent manner.
